# A platform for case-control matching enables association studies without genotype sharing

**DOI:** 10.1101/470450

**Authors:** Mykyta Artomov, Alexander A. Loboda, Maxim N. Artyomov, Mark J. Daly

**Author notes:** Authors declare no competing financial interests.

## Abstract

Acquiring a sufficiently powered cohort of control samples can be time consuming or, sometimes, impossible. Accordingly, an ability to leverage control samples that were already collected and sequenced elsewhere could dramatically improve power in all genetic association studies. However, since majority of the genotyped and sequenced human DNA samples to date are subject to strict data sharing regulations, large-scale sharing of, in particular, control samples is extremely challenging. Using insights from image recognition, we developed a method allowing selection of the best-matching controls in an external pool of samples that is compliant with personal genotype data protection restrictions. Our approach uses singular value decomposition of the matrix of case genotypes to rank controls in another study by similarity to cases. We demonstrate that this recovers an accurate case-control association analysis for both ultra-rare and common variants and implement and provide online access to a library of ~17,000 controls that enables association studies for case cohorts lacking control subjects.

## Introduction

Traditionally, genetic association studies require construction of a dataset consisting of both case and control genotypes. With this, one tries to eliminate all potential biases (technical or ancestral) between case and control cohorts, ensuring that discovered associations are phenotype-driven. While technical biases could be significantly diminished without explicit sharing of sensitive individual-level data by using the same sequencing technology and data processing standards for case and control cohorts, adjusting for population stratification requires researchers to either spend funds for sequencing study-specific controls (reducing the size of the case cohort) or file extensive paperwork to obtain what are sometimes a limited number of external control samples (roughly matched on ancestry and technology and consented for sharing) from public databases like dbGAP^1^. Moreover, re-using publicly available genetic data often requires each research team to process the data independently, performing redundant technical routines. The theoretical possibility of utilizing ancestry landscape for association studies without sharing genotypes was recently explored by UNICORN project^2^, proposing rigorous definition of a subject’s ancestry in a numerical way, but practical implementation of such concepts is lacking. Similarly, Guo *et al*^3^ recently proposed a methodology for usage of databases like gnomAD or ExAC as controls through calibration of association test statistics using benign variants, yet this approach is limited in ability to select matching control subjects for an association study.

In this work, we used insights from image recognition algorithms^4^ and developed a rigorous methodology for matching background variation in independent datasets without explicit genotype sharing. Specifically, we used singular value decomposition (**Supplementary Figure S1**) which expresses original matrix of genotypes as multiplication of three matrices:*G = USV^T^*; where U – is an orthogonal basis of principal directions (with the 1^st^ direction chosen along the largest variance in the original data, *S* – a diagonal matrix of singular values and *V* – is a matrix of principal components that provides coordinates of original samples in the basis of matrix *U*. Importantly, for a given genotype matrix *G* of cohort of cases, left singular vectors matrix - *U*, can be used alone to reconstruct original matrix of genotypes, and therefore can be shared unrestrictedly with remote database/control server since it does not contain any information on the individual level data, rather only providing information about variance directions in case cohort as a whole. We show that given several first vectors of matrix *U* derived from the case cohort data, one can project control genotypes on such case-derived basis and subsequently rank controls by their similarity to case cohort. Accordingly, optimal control cohort can be chosen by taking maximum number of controls that is still able to maintain null distribution of the association test statistic^5^.

We performed tests both on genotyping array data and large-scale exome sequencing dataset confirming successful control selection. Additionally, we matched same exome samples from 1000 Genomes project processed through different variant discovery pipelines in two different ways to a set of controls showing that given exome capture concordance joint processing is not required for returning sufficient set of matched controls. Finally, we illustrate method performance with rare-variant association study in early onset breast cancer cohort, performed without genotype sharing. We use the proposed approach to make freely accessible a set of 16,946 controls that can be used by researchers worldwide without sharing data restrictions. Specifically, we provide an R-package for case cohort data transformation (SVDFunctions) at the local machine and accompanying online portal called SVD-based Control Repository (SCoRe, www.dnascore.net), which selects a set of optimal controls and outputs corresponding control summary allele counts to enable association studies.

## Results

### Framework for control matching without genotype sharing

Following the standard workflow for genetic association studies – only autosomal LD-pruned DNA variants should be used for ancestral matching^6,7^. This ensures that downstream analysis reflects global LD structure rather than local LD. Case genotypes for common autosomal, LD-pruned variants could be represented as a genotype matrix (with entries “0”,”1”,”2” - corresponding to the number of alternative alleles in a genotype). With singular value decomposition (SVD) one can obtain an orthogonal basis of principal directions in cases (*U* matrix, **Figure 1**). Coordinates of principal directions (left singular vectors), especially a limited number of them, on their own cannot be used to recover individual level genotypes. They represent orthogonal directions within original data with the largest variance and do not contain any information of where original samples are located in this space. For reconstruction of the original matrix one needs to have some information about all three pieces of decomposition result^8^. From the matrix approximation properties of the SVD it is known that the first singular vector represents “dominating” direction of the data matrix. Thus, if folded back to case genotypes, vectors *ui* should be similar in global LD structure to the case cohort.

**Figure 1.**
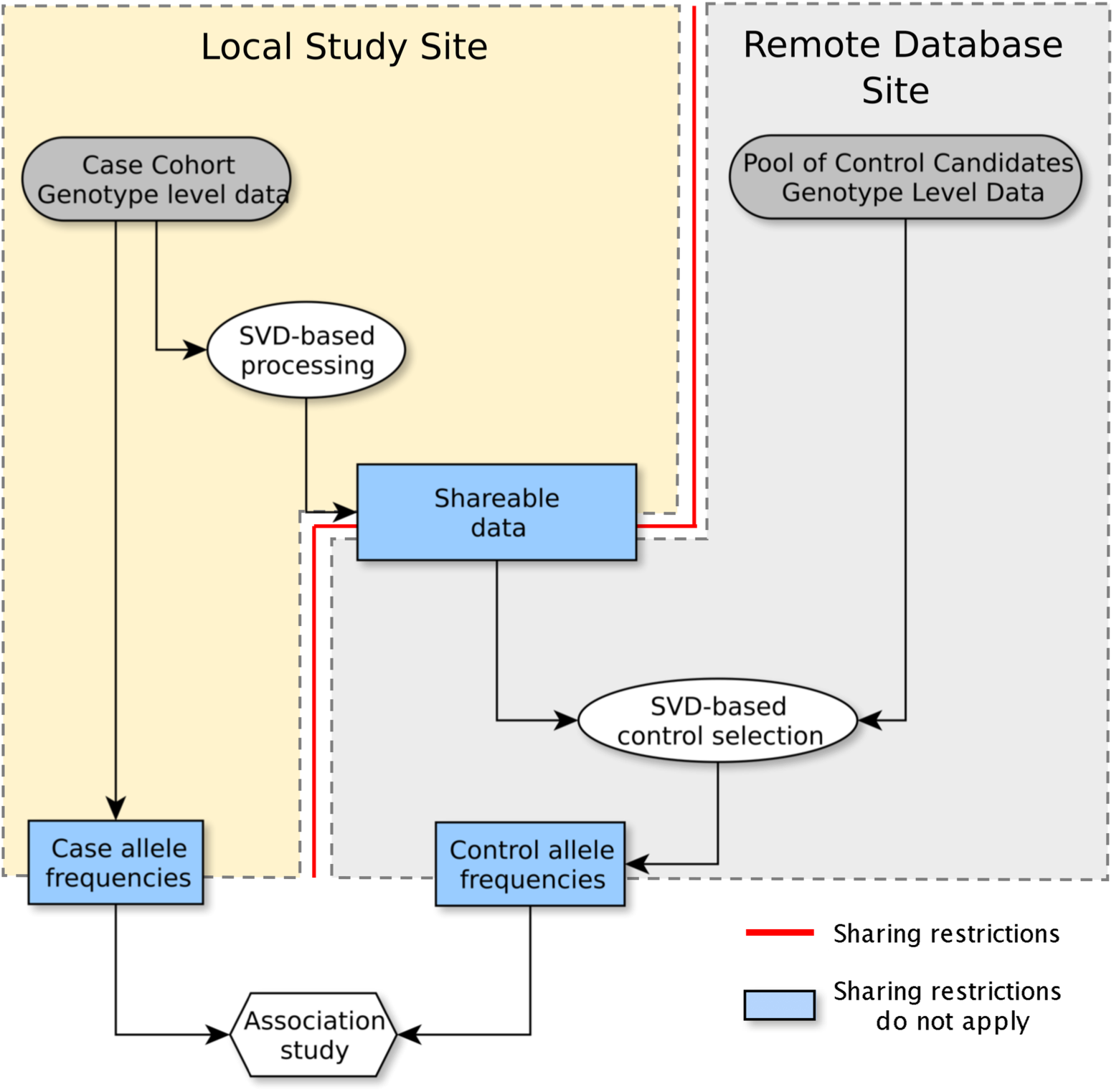
Scheme of an association study without genotype sharing. Individual level genotype data is subject to data sharing restrictions. SVD-based processing creates anonymous data describing variation in case genotypes without storing individual data that could be shared with no restrictions. Remote server with a pool of controls selects a set of controls genotype variation matching cases, estimates allele frequency for sites to be used for association study and delivers results to the user.

It is established that these methods perform well if the following conditions are met: each sample could be well characterized by a few of the first singular vectors; an expansion in terms of the first few singular vectors discriminates well between the sample ancestries^4^; cases represent a relatively homogenous ancestral cluster (that is, if you are conducting a meta-analysis across two diverse ancestries, you should run the two case groups separately). Therefore, we can compute how well a prospective control sample can be represented in the basis of case cohort. This can be done by computing residual vector in the least squares problem of the type:

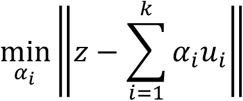

where *z –* a prospective control sample genotype vector, *ui –* left singular vectors of the case genotype matrix SVD, k – first few singular vectors. Norm of the residual vector in this case could be computed as ‖(I − UU^T^)Z‖_2_. SVD requires access to raw genotypes so must be completed locally and limited basis of the left singular vectors {*u*k} could be unrestrictedly shared. External samples that allow general research use could be stored on the remote control server (**Figure 1**) accepting anonymous SVD-processed data from local case clients. Remote control server then estimates residual norm for every prospective control and creates ranking for the representation quality in the basis of case cohort. Case allele frequencies need to be supplied to the control server for defining appropriate size of control set. For every control residual vector norm threshold an association test should be performed using allele frequencies for DNA variants used for matching and genomic inflation factor^5^ estimated. Largest set of controls delivering acceptably null distribution of the test statistic (λ_GC_≤1.05) should be then taken as a matched set of controls. Allele frequencies for variants of interest should then be computed in matched control dataset and could be shared with client to run full-scale association test (Toy example of case-control study with 2 cases and 2 controls is explained at **Supplementary Figure S2**). Unlike individual level data, such summary statistic sharing from most consented resources is routinely allowed^9,10^.

### 1000 genomes. OMNI array genotyping data

To validate our approach, we used 1000 genomes genotyping data. Entire dataset was subjected to per genotype and per individual quality check procedures (**Supplementary Figure S3**). Final dataset consisted of 10,000 LD-pruned autosomal variants and 1,708 non-related samples (**Figure 2A**). To simulate an association study we divided the dataset into 100 “cases” of European ancestry (as provided by 1000 genomes project annotation) and “control pool” of 1608 samples that included 359 European samples (**Figure 2A,B**). We simulated an association study without genotype sharing by separating every case cohort from control candidates (**Figure 2C**). For a case group SVD was computed and top 5 left singular vectors were used for control matching purposes. Residual vector norm was computed for every control, ranking controls by similarity to case cohort (**Figure 2D**) and separating ancestries (**Figure 2E**). Choosing different residual norm thresholds allows to create different pools of controls of variable matching quality (**Figure 2D,E**). Indeed, PCA plot built using shared genotypes confirmed that increase in residual vector norm threshold results in departure from original European case cluster delivering poor control matching quality (**Figure 2F**). To select optimal threshold value, allele frequency of variants in cases is used to estimate association test statistics and genomic control factor for different residual vector norm thresholds (**Figure 2G,H**). Largest control pool size with null distributed association test statistics (e.g. λ_GC_≤1.05) was selected as optimally matched set (**Figure 2I**). 100 simulation instances with random 100 “case” cohorts (100 European samples each) yielded on average selection of 428 controls (SD=26.6) with estimated mean genomic inflation λ_GC_=1.049 (SD=8.76×10^-4^) (**Figure 2J**). These control sets covered nearly all European samples in Control Pool and also selected a small number of controls annotated in 1000 Genomes as Latin-Americans (though still keeping genomic inflation factor below 1.05). In 100 random selections of “case” cohort (100 European samples each), we identified a group of Latin-Americans recurrently selected for a matched control set (**Supplementary Figure S4A**). It appears to be the closest to Europeans on PCA (**Supplementary Figure S4B**). Selection of samples with non-matching ancestry annotation is bound to the statistical power of the association testing used for genomic inflation estimation, thus, the more cases are used for association testing the fewer controls of non-matching ancestry annotation are selected (**Supplementary Figure S4C**). Though, with any set of case cohort genomic inflation factor is below 1.05 implying neglectable effect on further phenotypic association test results (**Supplementary Figure S4D**).

**Figure 2.**
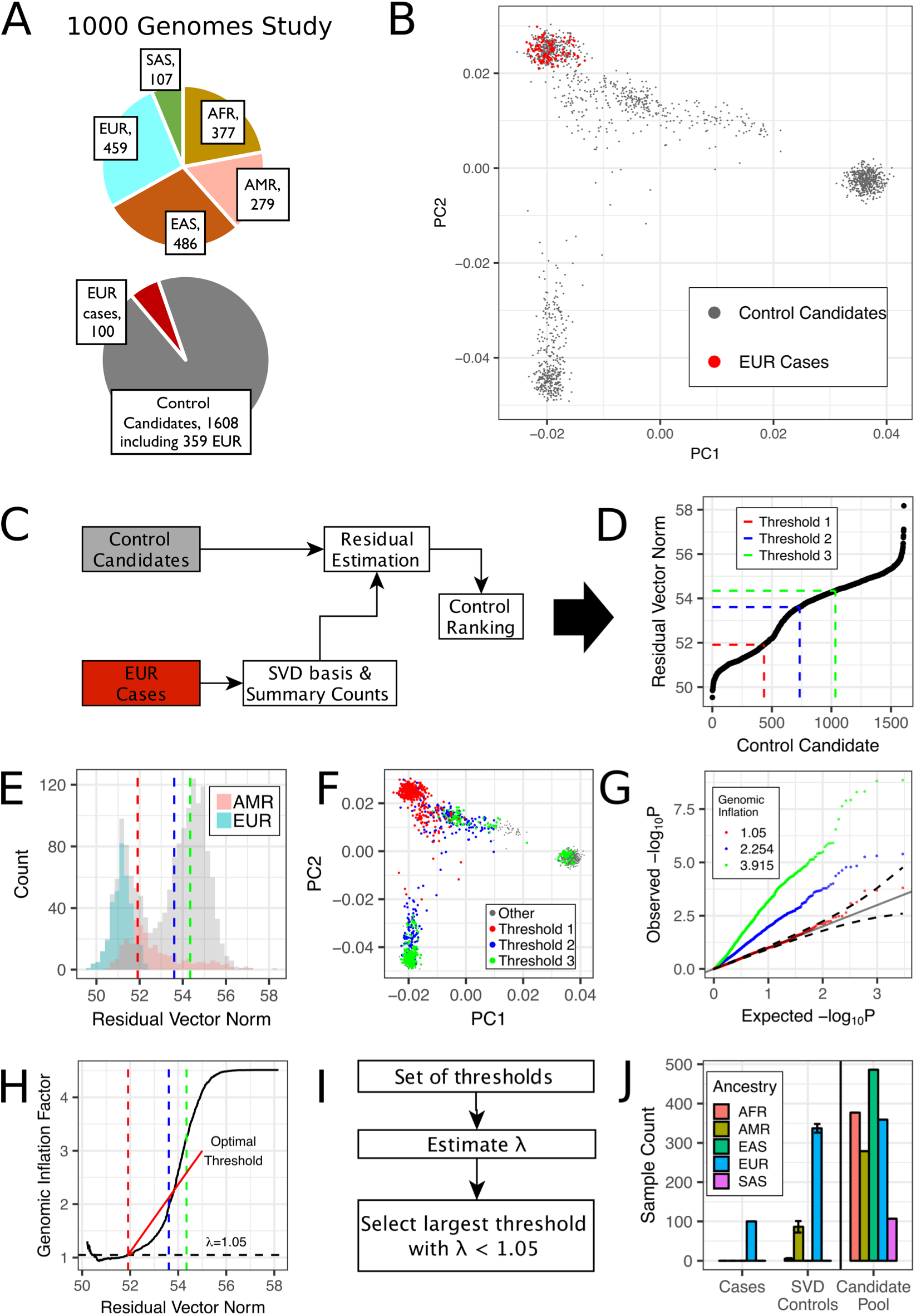
Case Study 1: Simulated Case-Control Study with 1000 Genomes OMNI Genotyping Array Data. (A) Breakdown of ancestries present in the dataset and case-control study set up: random 100 European samples are selected as “cases” and the rest of the data is tested as prospective controls; (B) Conventional PCA showing European samples selected as case group; (C) Scheme of data handling simulating association study without genotype sharing; (D) Ranking of controls by quality of representation in case basis; (E) Distribution of residual vector norms for prospective controls; (F) Conventional PCA shows that greater values of residual vector norms correspond to greater departure from European cluster; (G) Quantile-Quantile plots for multiple thresholds demonstrate increasing inflation; (H) Optimal residual vector norm threshold should deliver λ_GC_<1.05; (I) Optimal threshold selection scheme; (J) Matching experiment summary results.

Furthermore, such SVD-based approach can be used for creating an easy-to-use ancestry predictor that yield large confidence results (**Supplementary Methods, Supplementary Figure S5**, SVDFunctions R-package available at https://github.com/alexloboda/SVDFunctions).

### Exome sequencing data

Further, we sought to test how this approach performs in exome sequencing data with relatively small number of LD-pruned DNA variants. We used an aggregated set of whole exome sequences consented for joint variant calling resulting in 37,607 samples to build a test dataset (**Supplementary Table S1**). Further, a subset of LD-pruned variants used for PCA in ExAC database^9^ was subjected to quality check: only genotypes with DP>10, GQ>20 and variants with less than 500 missing genotypes were allowed, resulting in 4,561 DNA variant left for analysis. Samples were further analyzed for relatedness and related samples were removed: in pairs of samples with π̂ < 0.2 only one sample was randomly kept (**Supplementary Methods**). Final dataset consisted of 32,677 whole exome sequences (**Figure 3A**). Similarly, to the experiment with genotyping data that was described above, we randomly selected 3,000 European samples as a “case” set and leftover 29,677 samples became a disjoined pool of controls (**Figure 3B**). We used first 5 singular vectors to estimate residual norm for every sample in the control set and create a control ranking (**Figure 3C**), which distinctly separated ancestral groups (**Figure 3D**). Increase in residual norm results in departure from targeted European control cluster and inflation of the association test statistic (**Figure 3E,F**). Optimal control set size was defined as a largest set of controls with genomic inflation factor less or equal to 1.05 or the smallest inflation factor if no values below 1.05 are available (**Figure 3G**). In 100 random selections of “case” cohort (3000 Europeans each) on mean control set size was 17,435 (SD=921.97) with genomic inflation factor mean λ_GC_=1.056 (SD=0.012) (**Figure 3H**).

**Figure 3.**
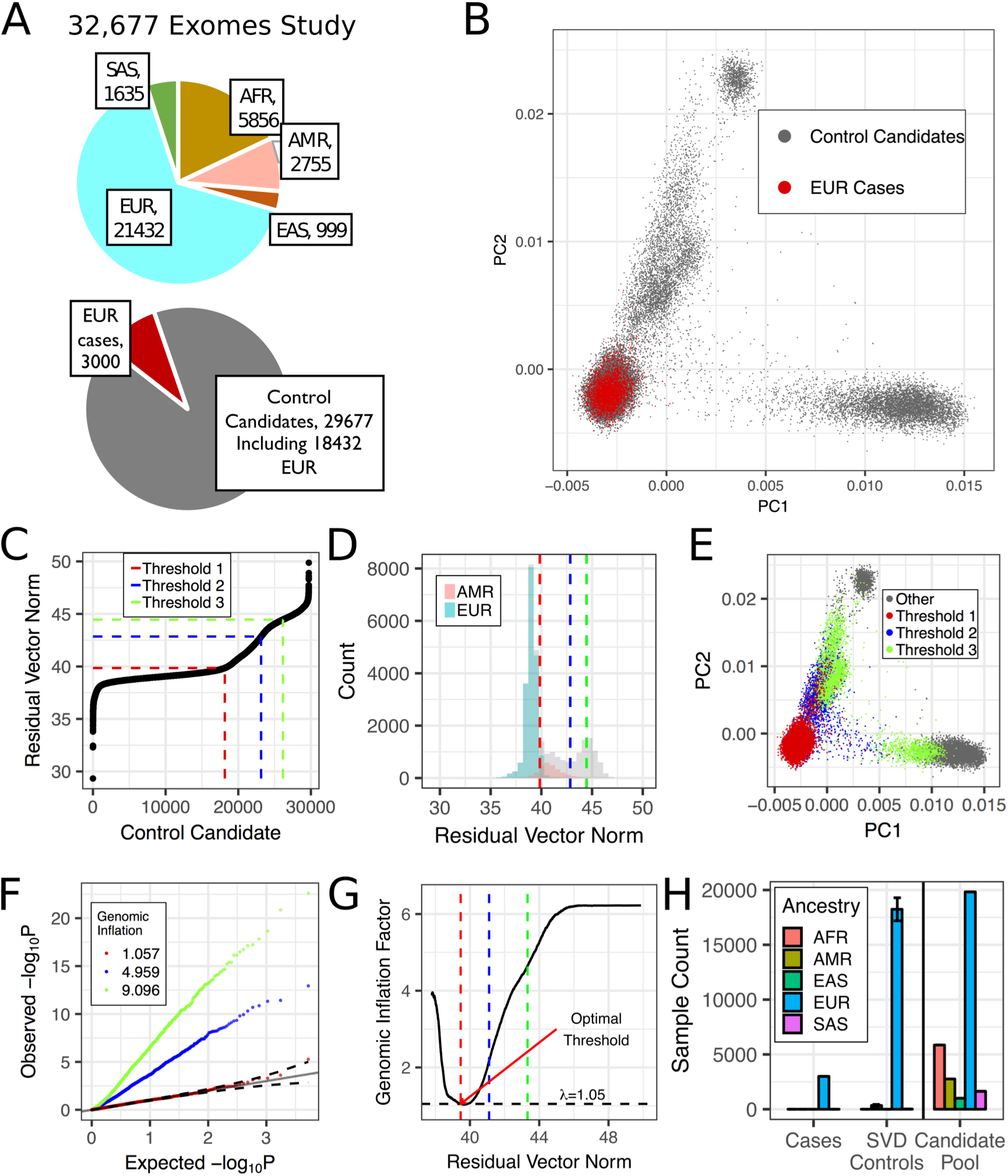
Case Study 2: Simulated Case-Control Study with 32,677 Exomes. (A) Breakdown of ancestries present in the dataset and case-control study set up: random 3000 European samples are selected as “cases” and the rest of the data is tested as prospective controls; (B) Conventional PCA showing European samples selected as case group; (C) Ranking of controls by quality of representation in case basis; (D) Distribution of residual vector norms for prospective controls; (E) Conventional PCA shows that greater values of residual vector norms correspond to greater departure from European cluster; (F) Quantile-Quantile plots for multiple thresholds demonstrate increasing inflation; (G) Optimal residual vector norm threshold should deliver λ_GC_<1.05; (H) Matching experiment summary results.

We analyzed how number of selected for analysis singular vectors affects size of matched controls cohort. Upon increase in number of singular vectors, more variance could be explained in limited reconstruction of the original matrix (e.g. if both *U, S, V* matrices are available for original genotype matrix, its reconstruction using first several vectors of *U* matrix would capture the variance better if more vectors are used, **Supplementary Figure S6A,B**). Though, since in our algorithm no matrix reconstruction is performed, there is no significant dependence of matched controls cohort size on the size of the case-derived basis (**Supplementary Figure S6C,D**). Thus, it is reasonable to use only first few singular vectors to increase computation speed.

### Controlling for technical bias

In the above examples, genotyping or sequencing data was processed in the uniform way and in case of exomes – variants were called jointly, an action which is impossible without explicit genotype sharing. Thus, we looked into effects of technical biases on the remote case-control matching. We used publicly available 1000 genomes Phase 3 data^10^ to select 46 samples (CEU ancestry), that were also present in our exome control pool dataset and attempted to match them to a control group. For this analysis control pool dataset was modified to exclude all 1000 genomes samples. Samples with known cancer phenotype were also excluded for further use of this data as a public database of controls (**Supplementary Figure S7A**). Both “external” and jointly processed 1000 genomes samples were acceptably matched to a group of controls (**Supplementary Figure S7B-D**), with exactly the same 5963 control candidates selected for both case groups and 1 additional control matched to “internal” cases. Important to note, that external 1000 genomes exomes were sequenced using similar to control pool’s Agilent exome capture kit. Therefore, we conclude that separate variant calling does not introduce major technical biases within the proposed approach.

One the level of experimental design, difference in DNA sequence capture used for exome sequencing are known to introduce major effects in variant calling and cannot be cross-used for association tests even if variants are jointly called^9,11^. The proposed approach will automatically treat platform-based bias as a kind of “ancestry”-matching and control candidates from the same sequencing platform would be prioritized over other options. However, in this initial release of 17,000 we only provide the control samples sequenced with Whole Exome Agilent 1.1 RefSeq plus 3 boosters capture. Being the most represented capture in ExAC (~77%)^9^ covers a major part of sequenced samples to date.

Additional data quality metrics: depth, missing data rate could be directly shared between control server and case client. Even with direct sharing of genotypes further work needs to be done to get a gold-standard association study: lining up depth and accuracy at every site, gene, exon. While control platform described here will ensure ancestral and platform matching it is up to a researcher to match quality metrics and conduct careful statistical analysis.

### Breast Cancer Cohort Analysis

We implemented presented methodology in combination of R-package (SVDFunctions) for data preprocessing and online SVD-based Control Repository (SCoRe) platform (www.dnascore.net) and performed a gene-based association study for a cohort of early onset breast cancer patients (dbGAP: phs000822.v1.p1): 291 non-related cases matching quality standards were further used for analysis. Genotype matrix and summary genotype counts of cases were constructed for 4,561 LD-pruned DNA variants that are available for matching through SCoRe. 1,930 controls were matched to the case cohort with λ_GC_=1.07 (**Figure 4A**). To ensure that not only the targeted DNA variants are matched we inquired SCoRe to return summary allele counts for various variant classes found in cases: common synonymous variants (that were not used for matching), rare synonymous variants grouped by gene (singletons in cases, implying MAF<0.00172). Null distribution of the test statistic in both inquiries confirmed case-control matching in both common and rare variant background variation (**Figure 4D,E**). Further, rare (MAF<0.00172) protein-truncating variants (PTV) gene burden was examined. 1930 genes had at least 1 PTV variant in cases (Bonferroni corrected significance threshold 0.05/1930 = 2.56×10^-5^). These genes were submitted to SCoRe to obtain summary gene-based counts of PTVs in matched controls and further, association test was performed locally using Fisher’s exact test (**Figure 4F**) re-discovering *BRCA1* as a susceptibility gene (with 12 carriers in 291 case and 7 carriers in 1930 controls; Fisher test P=6.36×10^-7^).

**Figure 4.**
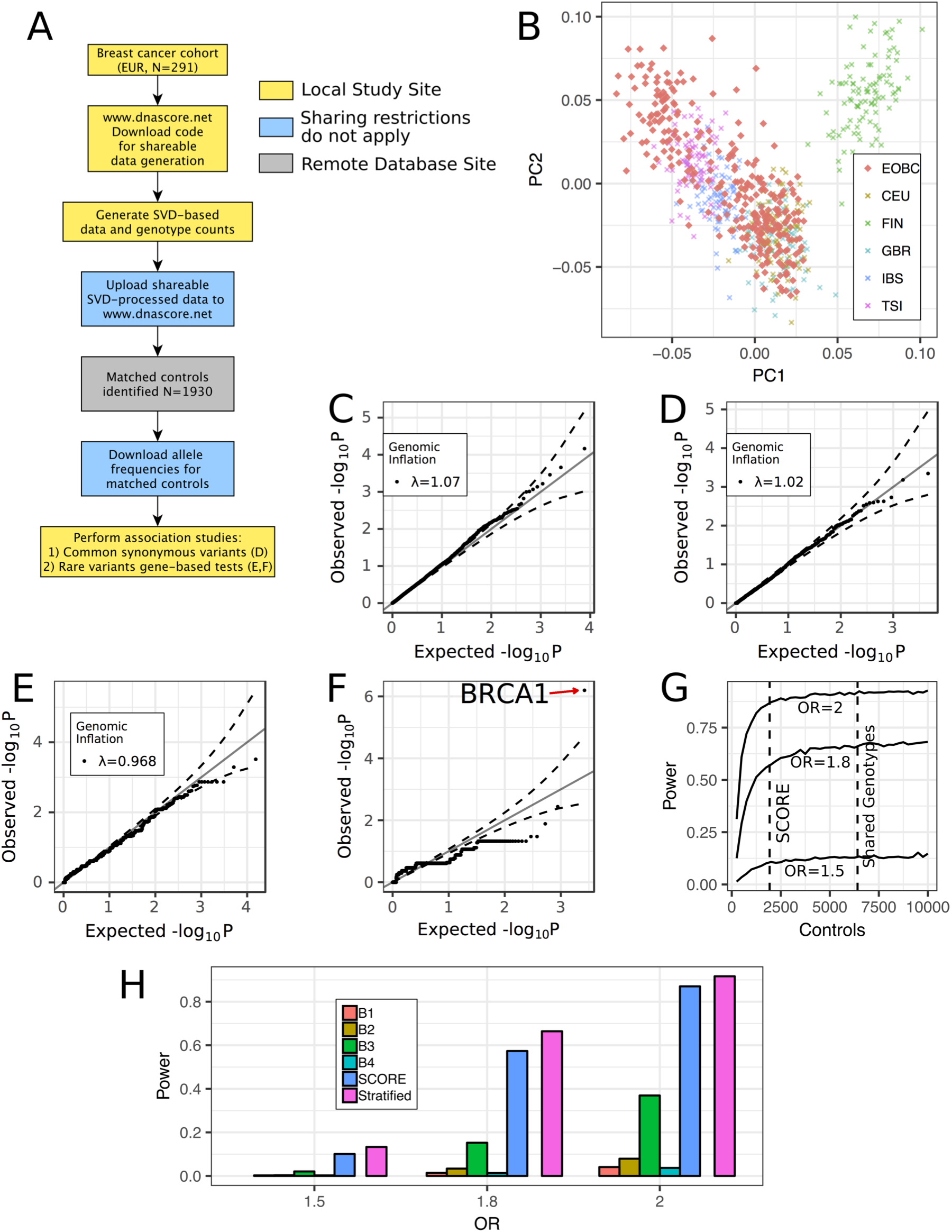
Case Study 3: Early-onset Breast Cancer Association Study. (A) Experimental workflow for obtaining matched controls data from SCoRe online platform; (B) Principal component analysis of breast cancer cohort (EOBC) performed jointly with 1000 genomes European samples. Multiple European sub-ancestries are present in case cohort, including substantial Ashkenazi group; (C) QQ-plot obtained from SCoRe platform for 1930 matched controls; (D) QQ-plot of association study performed on common (MAF>5%) synonymous variants found in cases with control allele frequencies obtained from SCoRe; (E) QQ-plot for gene-burden test for rare synonymous (singletons in cases, corresponds to MAF<0.00172) with gene-level summary allele counts obtained from SCoRe (F) QQ-plot for rare protein-truncating variants (singletons in cases, corresponds to MAF<0.00172) with gene-level summary allele counts obtained from SCoRe. BRCA1 is the top associated gene; (G) Fisher test statistical power estimate for 291 cases. Control cohorts matched with SCoRe platform and conventional stratified analysis with genotype sharing (**Supplementary Figure S8**) are labeled with dashed lines; (H) Fisher test statistical power for each case-control cohort: separate batches from shared genotypes analysis, stratified analysis with shared genotypes and SCoRe platform (**Supplementary Figure S8**).

Strikingly, despite the complex ancestral structure of case cohort with substantial amount of Ashkenazi samples (**Figure 4B**), we identified a set of controls that were matched as a single batch with λ_GC_=1.07 (**Figure 4C**). This was very different when compared to conventional genotype sharing association study: due to complex ancestral structure in cases, matching of controls as a single batch is largely impeded. We used K-means clustering to identify matching batches (**Supplementary Figure S8A, B**) so that all of them return null-distributed test statistic (using same variants as were used for SCoRe-based matching, **Supplementary Figure S8C-F**). Using batch-based matching with fully shared genotypes, we identified a total of 6,415 controls (out of 16,946 candidate controls in SCoRe) to the same case group in jointly called dataset.

Statistical power was estimated for Fisher’s exact test using multiple odds ratios for SCORE and conventional genotype sharing tests (**Figure 4G**), implying 291 case cohort and single-batch matched control cohorts of 1930 and 6415 samples, respectively (with disease prevalence 12.4%^12^). We also estimated power for separate batches in a non-stratified analysis set up (**Figure 4H**). The results show that SCoRe performs significantly better in a single-batch analysis than a shared genotype study and is producing sufficient number of controls to effectively saturate a power of the study in terms of size of control cohort.

### Web-portal for Case-Control Matching

We established a public online platform with 16,946 non-cancer controls available for matching (**Supplementary Table S2**). We provide a list of variants available for matching in control pool, and a R-package(SVDFunctions) for local generating of the case genotype matrix from standard PLINK^13^ binary file format and a code to generate SVD and allele counts data that could be uploaded to the matching portal. Available at www.dnascore.net.

We set a hard threshold on minimal number of matched controls (500 samples) to deliver allele frequencies to a user and controls are selected in batches of 10. This ensures only anonymous summary allele frequencies are disclosed from control cohort. Quires to the database could be asking for variant based frequencies or cumulative counts of rare variants per gene. For variant-based quires we limit return to common variants (MAF>1%) to avoid disclosing unique DNA variants that would readily disclose a genotype.

## Discussion

Statistical power is a key to successful association study and control set size is often a limiting factor. Despite potential availability of control sets through public repositories, large efforts should be put into processing case and control datasets jointly before even preliminary results of an association study could emerge. Practically, this often becomes infeasible for the small cohort studies limited to data access or computational power. Assembly of large case-control datasets is generally done by international consortia (ExAC, PGC, IBD Genetics Consortium, etc.) as this requires a lot of effort and generous funding. We provide a large pool of exome sequences and a tool enabling rapid selection of matched control sets without genotype sharing that ultimately provides allele frequency statistics required for performing association tests. Nearly no effort is required from the user side to get all information needed for association study, facilitating future discovery of associated genes and DNA variants.

Local cohorts assembled at hospitals as a part of clinical screening procedures or genetic counselling often have very modestly sized or none control sets and stringent sharing regulation. Our platform enables case-control study design and boosts statistical power for such patient cohorts. Especially, for rare mendelian phenotypes where assembly of well-powered case-control cohort is impeded by low disease prevalence.

Finally, hundreds of thousands samples were subjected to exome or genome sequencing to date in the world. However, all this data exists in isolated pieces limiting potential benefit for genetic studies. We provide a repository of the software codes used for running the matching platform so that it could readily be implemented by large data holders – National biobank initiatives and international disease consortia to let community benefit from large-scale genetic resources. This is also critically important for advancing genetic association studies in situations when explicit data sharing is not permitted or very challenging in the international settings, thus potentially providing insights into rare sample collections that were not available so far. Finally, approach developed in this work creates a path to creating unified central repository that would encompass all studies published in dbGAP and make it accessible to association studies run in any design and cohort.

## Methods

### 1000 genomes OMNI-array genotyping data

Raw genotypes were downloaded from 1000 genomes FTP site at: ftp://ftp.1000genomes.ebi.ac.uk/vol1/ftp/release/20130502/supporting/hd_genotype_chip/ Dataset was subjected to variant and samples quality check, according to the standard genotyping data handling protocol (**Supplementary Figure S2**)^6,7^. Quality check was performed with PLINK package^13^. Code for converting binary PLINK format files into genotype matrix used for matching is available through a control matching web-portal at

### Exome Sequencing Data

Whole exome libraries were prepared using Whole Exome Agilent 1.1 RefSeq plus 3 boosters capture kit and protocol, automated on the Agilent Bravo and Hamilton Starlet. Libraries were then prepared for sequencing using a modified version of the manufacturer’s suggested protocol, automated on the Agilent Bravo and Hamilton Starlet, followed by sequencing on the Illumina HiSeq 2000. We used an aggregated set of samples consented for joint variant calling resulting in 37,607 samples (**Supplementary Table S1**). All samples were sequenced using the same capture reagents at the Broad Institute and aligned on the reference genome with BWA^14^ and the best-practices GATK/Picard Pipeline, followed by joint variant calling with all samples processed as a single batch using GATK v 3.1-144 Haplotype Caller^15–17^. The resulting dataset had 7,094,027 distinct variants. Variant effect predictor was used for variant annotation^18^.

Early onset breast cancer cohort used for association study is available through dbGAP (phs000822.v1.p1).

1000 genomes sequencing data was downloaded from ftp://ftp.1000genomes.ebi.ac.uk/vol1/ftp/release/20130502/

### Software

Software pipeline was developed using R-language and libraries^19–21^. PLINK/SEQ library was used for operations with sequencing data^22^.

Source code, usage examples and manual are available at supplemental materials and online: www.dnascore.net and https://github.com/alexloboda/SVDFunctions.

## Acknowledgements

Authors would like to thank Konstantin Zaitsev (Washington University in St. Louis) for helpful advice on singular value decomposition methodology. M.A. would like to thank Dr. Andrey Shaw (Genentech) for inspiration to pursue research in human genetics.

## Author Contributions

Conceptualization: M.A., M.N.A, M.J.D. Investigation: M.A., A.A.L., M.N.A., M.J.D. Software: M.A., A.A.L. Writing Original Draft: M.A., M.N.A., M.J.D.

## References

1. Mailman, M. D. et al. The NCBI dbGaP database of genotypes and phenotypes. Nat. Genet. 39, 1181–1186 (2007).

2. Bodea, C. A. et al. A Method to Exploit the Structure of Genetic Ancestry Space to Enhance Case-Control Studies. Am. J. Hum. Genet. 98, 857–868 (2016).

3. Guo, M. H., Plummer, L., Chan, Y.-M., Hirschhorn, J. N. & Lippincott, M. F. Burden Testing of Rare Variants Identified through Exome Sequencing via Publicly Available Control Data. Am. J. Hum. Genet. 103, 522–534 (2018).

4. Elden, L. Matrix methods in data mining and pattern recognition. Society of Industrial and Applied Mathematics. (2007).

5. Devlin, B. & Roeder, K. Genomic control for association studies. Biometrics 55, 997–1004 (1999).

6. Anderson, C. A. et al. Data quality control in genetic case-control association studies. Nat. Protoc. 5, 1564–1573 (2010).

7. Reed, E. et al. A guide to genome-wide association analysis and post-analytic interrogation. Stat. Med. 34, 3769–3792 (2015).

8. Abdi, H. Singular Value Decomposition (SVD) and Generalized Singular Value Decomposition (GSVD). in Encyclopedia of Measurement and Statistics 907–912 (Thousand Oaks (CA): Sage, 2007).

9. Lek, M. et al. Analysis of protein-coding genetic variation in 60,706 humans. Nature 536, 285–291 (2016).

10. Consortium, T. 1000 G. P. A global reference for human genetic variation. Nature 526, 68–74 (2015).

11. Shigemizu, D. et al. Performance comparison of four commercial human whole-exome capture platforms. Sci. Rep. 5, 12742 (2015).

12. Cancer Statistics Facts. National Cancer Institute. (2018).

13. Purcell, S. et al. PLINK: A Tool Set for Whole-Genome Association and Population-Based Linkage Analyses. Am. J. Hum. Genet. 81, 559–575 (2007).

14. Li, H. & Durbin, R. Fast and accurate short read alignment with Burrows-Wheeler transform. Bioinformatics 25, 1754–1760 (2009).

15. DePristo, M. A. et al. A framework for variation discovery and genotyping using next-generation DNA sequencing data. Nat. Genet. 43, 491–8 (2011).

16. McKenna, A. et al. The Genome Analysis Toolkit: a MapReduce framework for analyzing next-generation DNA sequencing data. Genome Res. 20, 1297–303 (2010).

17. Van der Auwera, G. A. et al. From FastQ data to high confidence variant calls: the Genome Analysis Toolkit best practices pipeline. Curr. Protoc. Bioinforma. 43, 11.10.1-33 (2013).

18. McLaren, W. et al. The Ensembl Variant Effect Predictor. Genome Biol. 17, 122 (2016).

19. Team, R. C. A language and environment for statistical computing. (2013).

20. Wickham, H. Elegant Graphics for Data Analysis. (Springer-Verlag, 2016).

21. Clayton, D. snpStats: SnpMatrix and XSnpMatrix classes and methods. R package version 1.30.0. (2017).

22. Https://atgu.mgh.harvard.edu/plinkseq/.PLINK/SEQ.

